# The *sev-GAL4* driver in *Drosophila melanogaster* does not express in the eight pairs of dorso-medial and some other neurons in larval ventral ganglia: A correction

**DOI:** 10.1101/2024.12.05.627116

**Authors:** Vanshika Kaushik, Subhash C. Lakhotia

## Abstract

The *sev-Gal4* driver is widely used in *Drosophila* to express the target gene in specific subsets of cells in ommatidial units of developing eye. A 2015 report (Ray & Lakhotia, J Genet 94, 407-416) from our laboratory claimed that besides the eye disc cells, the *Sev-Gal4* (Bloomington stock 5793) also expresses in 8 pairs of dorso-medial neurons and some other cells in larval and early pupal ventral ganglia. The current study reveals that this claim was incorrect since the *UAS-GFP* transgene in Bloomington stock 1521 used as reporter in the 2015 study expresses in the 8 pairs of dorso-medial neurons and some other cells in larval and early pupal ventral ganglia even in undriven condition. The *UAS-eGFP* reporter in the BL-5431 stock, however, does not express in these ganglia, neither in undriven nor in *sev-Gal4* driven condition. This was also confirmed by the G-TRACE cell lineage study. Present results suggest that only four dorsal-midline cells in ventral ganglia and a cluster of cells in central region of brain hemisphere, besides the earlier known cells in eye disc and optic lobe of brain, express the *sev-Gal4* transgene in the stock 5793. The essentiality of examining undriven expression of a transgene cannot be under-emphasized.

## Introduction

Development of the *Gal4-UAS* system to express the desired gene in specific cell types at defined developmental stage (Brand and Perrimon 1993) is a remarkably versatile tool for understanding diverse aspects of developmental and induced gene expression. The *sevenless* (*sev*) gene, specifically expressed in R3, R4, R7 photoreceptors, cone cells and two mystery cells, is critical for development of ommatidial units in eye discs of third instar larvae (Banerjee *et al*. 1987; Tomlinson *et al*. 1987). The *sev-Gal4* driver has, therefore, been widely used to express the desired target gene, placed under the *UAS* promoter, to express in specific cells of developing eye discs. Since driving the expression of certain transgenes with *sev-Gal4* was reported to cause unexpected pupal lethality, our lab had earlier (Ray and Lakhotia 2015) examined the expression domains of the *sev-Gal4* transgene, using the *UAS-GFP* transgene and reported that besides the expected expression of the *sev-Gal4* in R3, R4, R7 photoreceptors, cone cells, and two mystery cells in developing ommatidia, this driver caused the reporter *UAS-GFP* transgene to express in certain other cell types as well. Prominent among these were 8 pairs of dorso-medial neurons in ventral ganglia of larval and early pupal stages. This work (Ray and Lakhotia 2015) also reported that another widely used eye-specific *GMR-Gal4* also expressed in the same set of dorso-medial neuron pairs in ventral ganglia. During our recent studies, however, we noticed that the expression domains of the *sev-Gal4* driver are different, specifically in the larval central nervous system, from our laboratory’s earlier report (Ray and Lakhotia 2015). We now note that the *UAS-GFP* reporter in the BL-1521 stock, used in the earlier study, shows an undriven leaky expression in the 8 pairs of dorso-medial neurons and several other cells in larval and early pupal ventral ganglia. Another *UAS-eGFP* reporter (BL-5431 form the Bloomington *Drosophila* Stock Center, USA), does not show any GFP expression in the earlier reported 8 pairs of dorso-medial ventral ganglia neurons, neither in undriven nor in *sev-Gal4*-driven background. Therefore, our lab’s earlier (Ray and Lakhotia 2015) report that the *sev-Gal4* (and *GMR-Gal4*) drivers express in 8 pairs of dorso-medial neurons and several other cells in larval and early pupal ventral ganglia was incorrect because undriven expression of the *UAS-GFP* reporter in the BL-1521 stock was not examined.

## Material and methods

### Fly stocks

The following fly stocks, maintained on standard agar cornmeal medium at 24^0^±1^0^C, were used in this study: i) *w*^*1118*^; *sev-Gal4* (Bloomington *Drosophila* Stock no. BL-5793, referred to here as *sev-Gal4*), ii) *w*^*1118*^; *UAS-GFP* (stock no. BL-1521, referred to here as *UAS-GFP*), iii) *w*^*1118*^; *UAS-eGFP* (stock no. BL-5431, referred to here as *UAS-eGFP*), and iv) *G-TRACE/ CyO* (stock no. BL-28280) (Pandey *et al*. 2024).The stocks i and ii are the same as used in the earlier study (Ray and Lakhotia 2015).

To drive expression of *UAS-GFP* or *UAS-eGFP*, the *w*^*1118*^; *sev-GAL4* flies were crossed with *w*^*1118*^; *UAS-GFP* or *UAS-eGFP* and the progeny larvae used for monitoring the GFP/eGFP reporter expression, respectively. To trace the lineage of *sev-Gal4* expressing neurons in the larval central nervous system, the progeny larvae of cross between *sev-GAL4* and *G-TRACE/CyO* flies were used. Synchronously developing larvae were obtained by collecting freshly hatched first instar larvae at 2 hr intervals, after discarding the first batch of hatched larvae. Eye discs and brain hemispheres with ventral ganglia (CNS) were dissected out from 96 hr old larvae (after hatching) in 1x PBS (1.37 M NaCl, 27 mM KCl, 100 mM Na_2_HPO_4_, and 20 mM KH_2_PO_4_, pH-7.4), fixed in 4% paraformaldehyde in 1xPBS for 20 min, and washed 3×15 min with 0.1% PBST (0.1% Triton X-100 in 1xPBS). The tissues were stained with DAPI (4′,6-diamidino-2-phenylindole,1µg/ml) for 20 min followed by 3×15 min washes with 0.1% PBST, and finally mounted in DABCO. The GFP expression was examined with a Zeiss 510 Meta laser scanning confocal microscope using the Zen software. All images were assembled using the Adobe Photoshop software.

## Results and Discussion

It has been reported (Ray and Lakhotia 2019) that sev-*Gal4* or *GMR-Gal4* driven expression of the *UAS-Ras*^*V12*^ transgene leads to substantial pupal death. To know if this is related to the reported (Ray and Lakhotia 2015) expression of *sev-Gal4* driven ectopic expression of activated Ras in the 8 pairs of dorso-median and some other neurons in ventral ganglia, we used the G-TRACE method (Evans *et al*. 2009; Pandey *et al*. 2024) to follow the lineage of these neurons. Surprisingly, no RFP or GFP expression (see Fig. 1) was detected in the 8 pairs of dorso-median and some other neurons in ventral ganglia of late 3^rd^ instar larvae that were claimed earlier (Ray and Lakhotia 2015) to express the *sev-Gal4* driver. Instead, four dorso-medially and 2-3 peripherally located neurons in ventral ganglia (Fig. 1a-c) and a cluster of 12-15 neurons in the mid-region of each of the brain hemispheres (Fig. 1d-f) showed GFP and/or RFP expression. It is notable that the RFP and GFP expression in different neurons was not always coincident (Fig. 1d-e), which may be due to variability in strength of the Gal4 driver and stability of RFP and GFP in different cells (Evans *et al*. 2009; Pandey *et al*. 2024).

**Fig 1.**
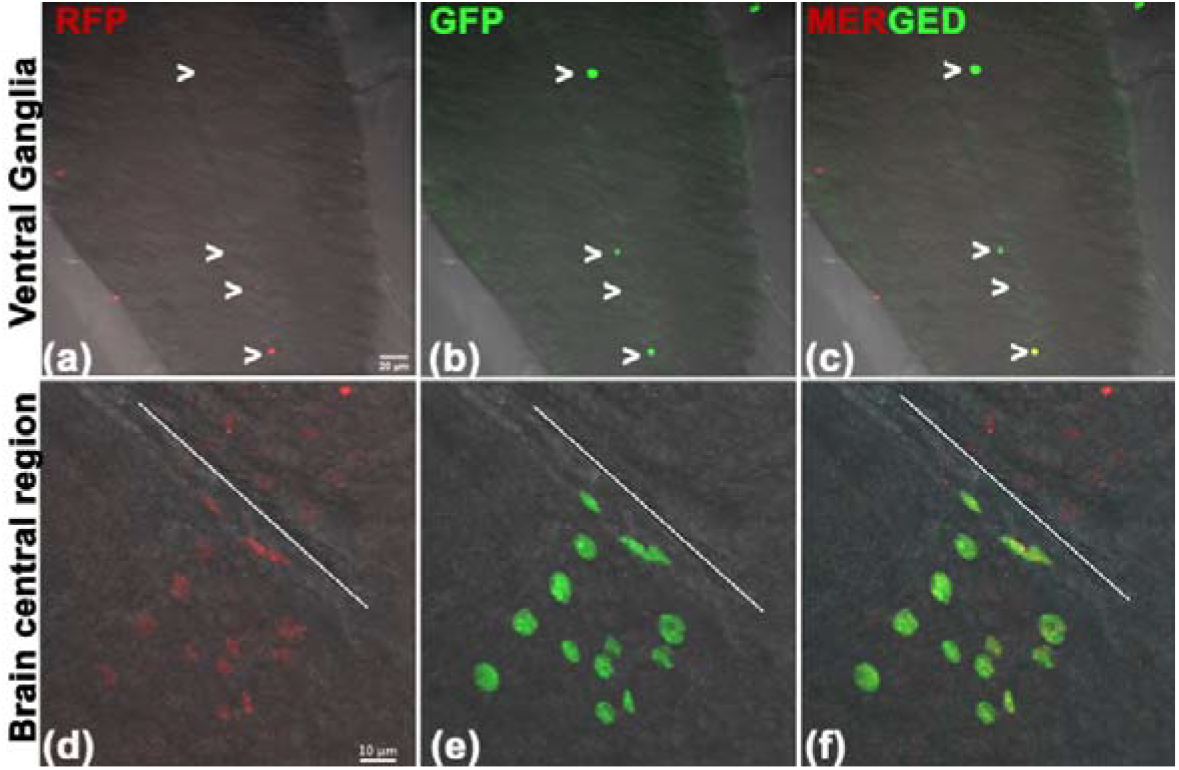
Confocal projection images showing expression of *sev-Gal4>G-TRACE* driven real-time RFP (red, **a, d**) and lineage GFP (green, **b, e**) fluorescence in the ventral ganglia (**a-c**) and central region of brain hemisphere (**d-f**) of late third instar larvae. White arrowheads in **a-c** indicate the RFP and/or GFP expressing neurons. The white line in **d-f** indicates the middle of the two cerebral hemispheres; note that a cluster of neurons shows RFP as well as GFP fluorescence on its left but on its right side, only faint RFP fluorescence is visible in similarly positioned cluster of neurons. Scale bar in **a** represents 20µm and applies to **a-c** while that in **d** represents 10µm and applies to **d-f**.

To resolve the absence of any G-TRACE lineage signal in the ventral ganglia neurons identified in the earlier report (Ray and Lakhotia 2015), we examined expression of two different GFP-reporters, viz., the *UAS-GFP* (BL-1521) and the *UAS-eGFP* (BL-5431) in un-driven and *sev-Gal4* driven conditions. As seen in Fig. 2a-2b’, the *UAS-GFP* reporter gene, used in our laboratory’s earlier study (Ray and Lakhotia 2015), expressed in the 8 pairs of dorso-median and several other neurons in the ventral region of ventral ganglia even in absence of the *sev-Gal4* driver (Fig. 2a, a’), all of which also showed GFP fluorescence in *sev-Gal4>UAS-GFP* larvae (Fig. 2b, b’). On the other hand, the *UAS-eGFP* reporter in BL-5431 stock did not express anywhere in ventral ganglia in undriven state (Fig. 2c, c’), but when driven by the *sev-Gal4* driver, eGFP-fluorescence was seen only in 4 dorso-medially placed neurons (white arrowheads in Fig. 2d, d’). Interestingly, ventral ganglia from *sev-Gal4>UAS-GFP* larvae also showed GFP fluorescence in the 4 dorso-medial neurons (white arrowheads in Fig. 2b) in addition to the undriven expression of GFP in the 8 pairs of dorso-median neurons and other cells in the ventral side (Fig. 2b, b’).

**Fig 2.**
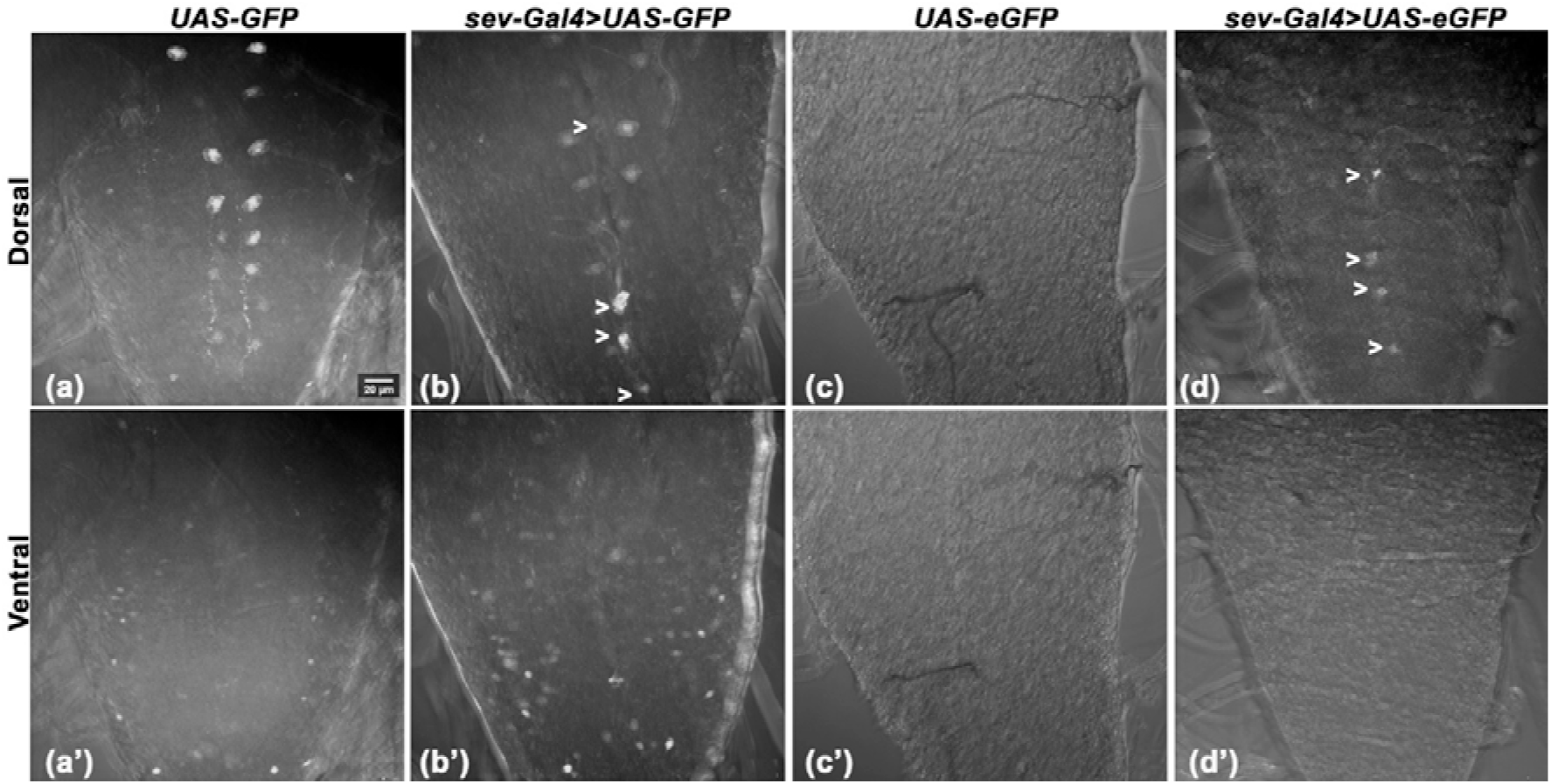
Confocal projection images of dorsal (**a-d**) and ventral (**a’-d’**) sides of the ventral ganglia of late third instar larvae showing nearly similar expression of *UAS-GFP* (white) in BL-1521 stock in 8 pairs of dorso-medial neurons (**a, b**) and many neurons on ventral side (**a’, b’**) in undriven (**a, a’**) as well as *sev-Gal4* driven (**b, b’**) conditions while the *UAS-eGFP* transgene in BL-5431 stock does not show GFP expression in any neurons neither in dorsal nor ventral sides of ventral ganglia in undriven condition (**c, c’**); in *sev-Gal4>UAS-eGFP* ventral ganglia (**d, d’**) only 4 dorso-median neurons (white arrowheads), but not the 8 pairs of dorso-median neurons, show GFP (white) expression. Scale bar in **a** represents 20µm and applies to all the images.

We next examined the undriven expression of these two GFP reporter lines in eye discs and brain regions (Fig. 3). While the undriven *UAS-GFP* transgene in BL-1521 did not express in any cell in eye disc (Fig. 3a), one cell in the optic lobe region of brain hemispheres (Fig. 3 b), a few neurons in the central part of the brain, closer to the midline, showed GFP fluorescence (Fig. 3c). The undriven *UAS-eGFP* reporter in the BL-5431 stock expressed only in one posteriorly located cell in the peripodial layer of eye disc (Fig. 3d) and two cells in optic lobe near the entry of optic nerve (Fig. 3e). The central region of brain did not show any undriven eGFP signal in the BL-5431 stock (Fig. 3f).

**Fig 3.**
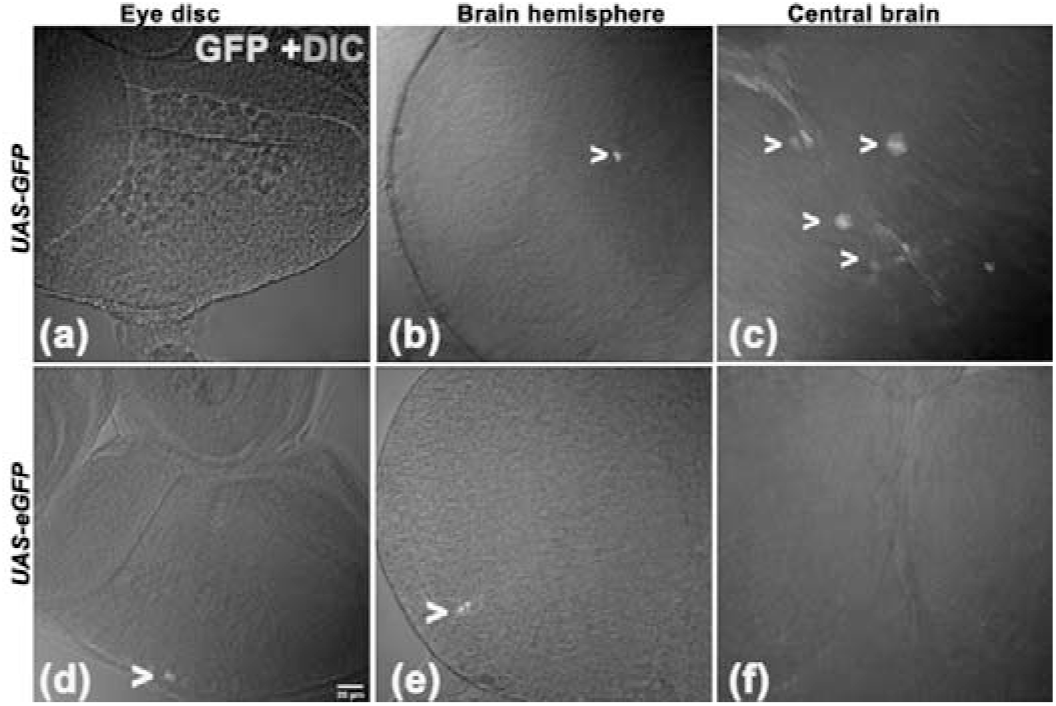
Single section confocal images of eye discs (**a, d**), brain hemisphere (**b, e**) and confocal projection images of central region of brain (**c, f**) of *UAS-GFP* (**a-c**) and *UAS-eGFP* (**d-f**) late 3^rd^ instar larvae. White arrowheads indicate cells that show undriven GFP (**b, c**) or eGFP (**d, e**) fluorescence. Scale bar in **d** represent 20µm and applies to all the images.

We compared the *sev-Gal4* driven expression of the *UAS-GFP* and *UAS-eGFP* transgenes in eye discs and optic lobes in the BL-1521 and BL-5431 stocks, respectively. As shown in Fig. 4, both the reporters exhibited identical fluorescence patterns that coincide with the known expression of Sevenless in developing ommatidia (Banerjee *et al*. 1987; Tomlinson *et al*. 1987). It is interesting that in both cases, the *sev-Gal4* driven GFP/eGFP fluorescence was also seen in the axons in the optic nerve (Fig. 4c, d). This indicates that contrary to the earlier belief (Ray and Lakhotia 2015) that the GFP localization following expression of the *UAS-GFP* transgene in the BL-1521 stock is nuclear, both the reporters (GFP/eGFP) can be nuclear as well as cytoplasmic and can move along the axons. In our experience, the eGFP reporter signal in the BL-5431 stock is weaker than the GFP reporter signal in the BL-1521 stock (Figs. 2-4).

**Fig 4.**
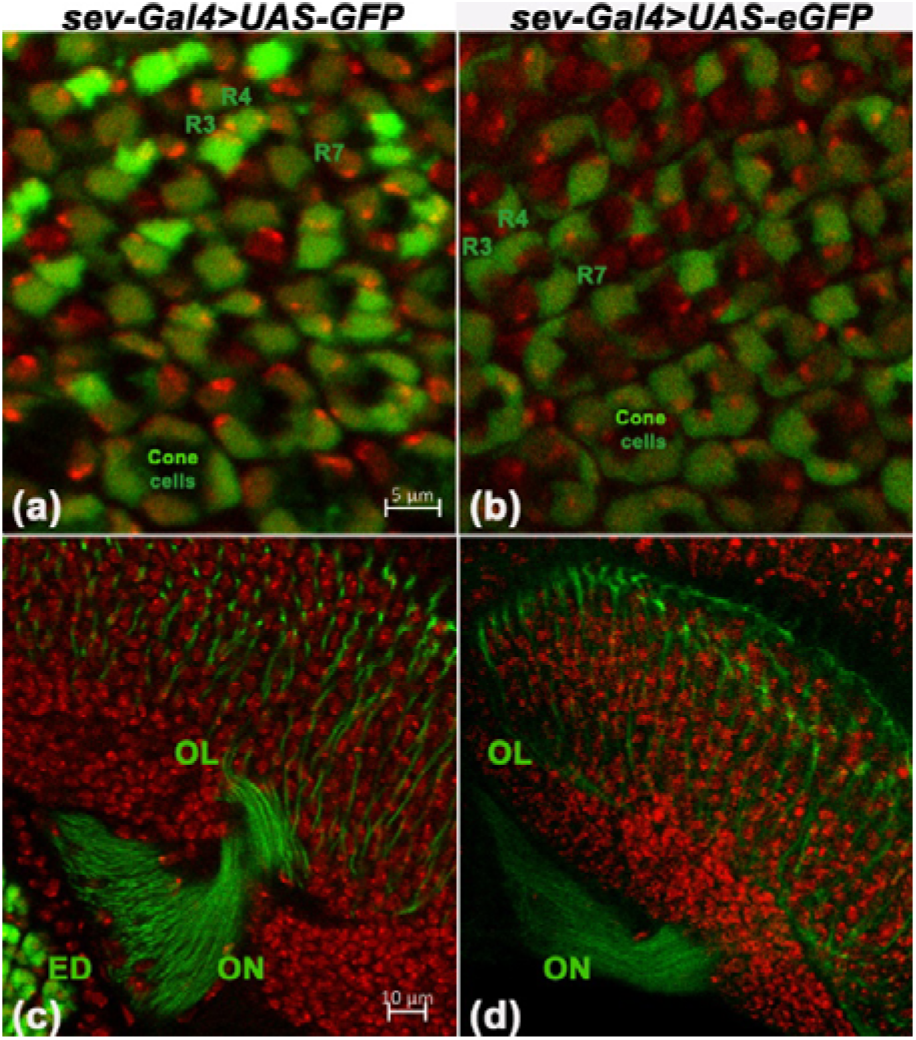
Single section confocal images of eye discs (**a, b**) and optic lobe regions of brain (**c, d**) from late third instar *sev-Gal4>UAS-GFP* (**a, c**) and *sev-Gal4>UAS-eGFP* (**b, d**) larvae showing GFP (green) and DAPI (red) fluorescence. The GFP expressing R3, R4, R7 photoreceptor and cone cells are marked in **a, b**. ED = eye disc, OL = optic lobe and ON = optic nerve. The scale bar in **a** represents 5µm and applies to **a-b**, that in **c** represents 10µm and applies to **c-d**.

Taken together, the present results of G-TRACE analysis and the GFP or eGFP expression in undriven and *sev-Gal4* driven conditions, show that the *sev-Gal4* driver is expressed in the CNS only in the 4 dorso-median neurons/cells in ventral ganglia, a cluster of neurons/cells in central region and a few other cells in brain hemispheres of late third instar larvae. The earlier claim (Ray and Lakhotia 2015) that the *sev-Gal4* driver expresses in 8 pairs of dorso-median and several other neurons in ventral side of larval ventral ganglia of *Drosophila melanogaster* was erroneous since the earlier study, unfortunately, did not examine this otherwise widely used reporter transgene’s undriven expression. It is notable that according to the information at the Bloomington *Drosophila* Stock Centre web site, the *UAS-GFP* transgene in the BL-1521 stock is inserted within the first intron of the *Myo31DF* gene. This makes it likely that this *UAS-GFP* transgene shows a ‘leaky’ expression in some cells due to its insertion within the *Myo31DF* host gene.

The lesson from our laboratory’s present and earlier (Ray and Lakhotia 2015) studies re-emphasizes the essentiality of checking the undriven ‘leaky’ expression of an inducible transgene, especially of a reporter gene under the *UAS* promoter to ensure that the expression domains under investigation are not erroneously inferred. The communicating author of this and the 2015 publication (Ray and Lakhotia 2015) regrets that the undriven expression of the *UAS-GFP* transgene in BL-1521 was not carefully examined in the 2015 study, which may have led to wrong inferences in some studies.

## Acknowledgements

We thank the Bloomington *Drosophila* Stock Center, USA, for fly stocks and the Department of Biotechnology, Government of India, New Delhi (BT/PR32126/BRB/10/1775/2019) for financial support.

## Notes

### Competing Interest Statement

The authors have declared no competing interest.

